# The fish body functions as an airfoil: surface pressures generate thrust during carangiform locomotion

**DOI:** 10.1101/2020.02.20.958389

**Authors:** Kelsey N. Lucas, George V. Lauder, Eric D. Tytell

## Abstract

The anterior body of many fishes is shaped like an airfoil turned on its side. With an oscillating angle to the swimming direction, such an airfoil experiences negative pressure due to both its shape and pitching movements. This negative pressure acts as thrust forces on the anterior body. Here, we apply a high-resolution, pressure-based approach to describe how two fishes, bluegill sunfish (*Lepomis macrochirus* Rafinesque) and brook trout (*Salvelinus fontinalis* Mitchill), swimming in the carangiform mode, the most common fish swimming mode, generate thrust on their anterior bodies using leading-edge suction mechanics, much like an airfoil. These mechanics contrast with those previously reported in lampreys – anguilliform swimmers – which produce thrust with negative pressure but do so through undulatory mechanics. The thrust produced on the anterior body of these carangiform swimmers through negative pressure comprises 28% of the total thrust produced over the body and caudal fin, substantially decreasing the net drag on the anterior body. On the posterior region, subtle differences in body shape and kinematics allow trout to produce more thrust than bluegill, suggesting that they may swim more effectively. Despite the large phylogenetic distance between these species, and differences near the tail, the pressure profiles around the anterior body are similar. We suggest that such airfoil-like mechanics are highly efficient, because they require very little movement and therefore relatively little active muscular energy, and may be used by a wide range of fishes since many species have appropriately-shaped bodies.

**Significance Statement:** Many fishes have bodies shaped like a low-drag airfoil, with a rounded leading edge and a smoothly tapered trailing region, and move like an airfoil pitching at a small angle. This shape reduces drag but its significance for thrust production by fishes has not been investigated experimentally. By quantifying body surface pressures and forces during swimming, we find that the anterior body shape and movement allows fishes to produce thrust in the same way as an oscillating airfoil. This work helps us to understand how the streamlined body shape of fishes contributes, not only to reducing drag, but also directly to propulsion, and, by quantitatively linking form and function, leads to a more complete understanding fish evolution and ecology.

## Introduction

It has long been appreciated that the shape of many fishes resembles a streamlined body (1–4). In particular, the two-dimensional horizontal cross-section through many fishes is similar in shape to modern airfoil profiles designed to minimize drag (3). Because nearly all aspects of a fish’s life depend on how well it swims, it has been suggested that this shape represents an evolutionary optimization to minimize drag for economical swimming (1). In general, swimming performance is linked to the evolution of fish body forms and movement patterns (5–10). For fishes that swim fast or migrate long distances, even small energy savings may be important.

However, along with reducing drag, an airfoil can directly generate propulsive forces by virtue of their shape and an effect called leading-edge suction. Due to its shape, an airfoil will generate a positive (above ambient) pressure stagnation point near its leading edge as flow divides to move along either side of the foil (3, 4, 11, 12). Then, the airfoil generates negative (below ambient) pressure over much of its length (Fig. 1B, and similar to the time-averaged pressure in Fig. 1A) (3, 4, 11–13). Since pressure produces a force perpendicular to the surface, negative pressure along the leading portion of the foil (∼5 to 40% in Fig. 1A,B) will contribute to thrust because the surface there is angled forward (illustrated in Fig. 1B) (4, 11, 14). Airfoils also produce thrust on their anterior regions through leading-edge suction when they are at an angle to the flow (12, 15, 16). When the airfoil is angled, the stagnation point and region of positive pressure is not directly on the tip of the airfoil (Fig. 1A,C) (15, 16). When the positive pressure deflects to one side, negative pressure moves forward to act more anteriorly on the opposite side (compare Fig. 1B,C) (15, 16). This area of negative pressure, positioned alongside forward-facing surfaces near the airfoil’s leading edge, acts as local forces with small thrust components in a mechanism called leading-edge suction (Fig. 1C) (11, 12, 14–16).

**Fig. 1.**
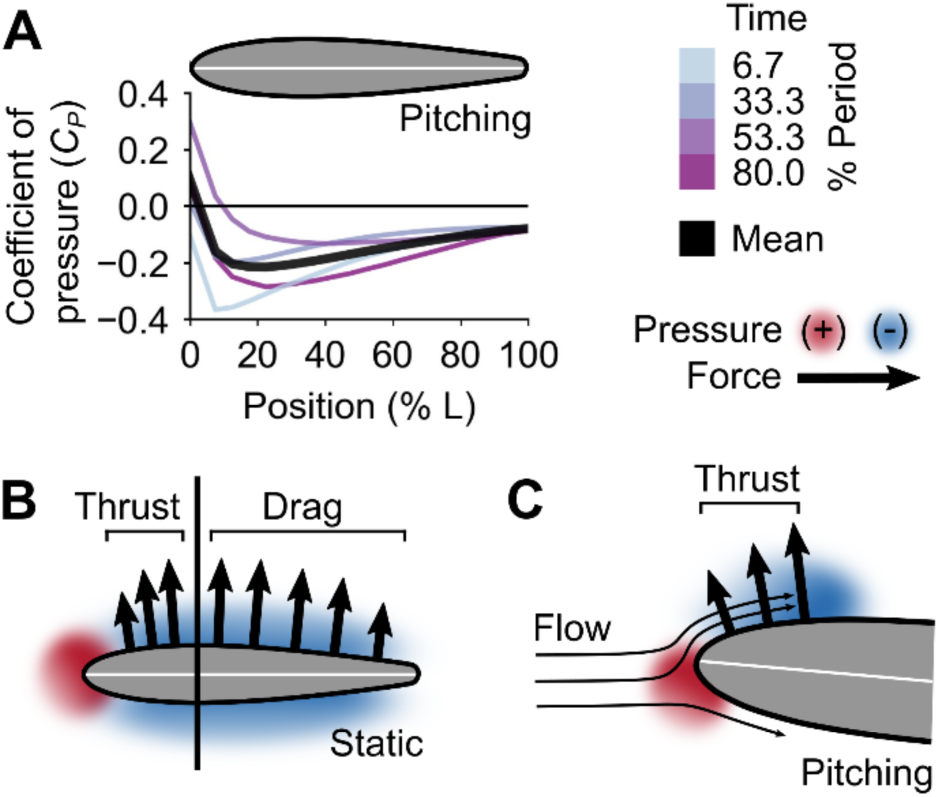
Physical mechanics of airfoils. (A) Coefficient of pressure (*C*_*P*_) along one side of a NACA 0015 airfoil with a rounded trailing edge pitching at reduced frequency 0.2, about 0° mean angle of attack, with an amplitude of ±5° (13). Colors indicate instantaneous pressure profiles, while the thick black line represents the time-averaged mean. (B) Pressure gradients around an airfoil (here, static at 0° angle of attack) act perpendicularly to the surface and can contribute to thrust or drag forces based on the orientation of the surface. (C) Leading-edge suction occurs when pitching movements of the airfoil shift the stagnation point and positive pressure to one side, allowing negative pressure to act more anteriorly on the opposite side (11, 12, 14–16). For clarity, in (B) and (C), only negative pressure forces on one side of the airfoil are shown.

If a fish’s body resembles an airfoil turned on its side, then we might expect that the anterior body might similarly produce thrust due to its shape and movements. Fishes that swim by primarily undulating the posterior half or less of their bodies in a range of patterns broadly classified as “carangiform” characteristically have airfoil-like bodies. But, while it has long been recognized that the airfoil-like shape of a carangiform swimmer is crucial for drag reduction (1, 4, 14, 17, 18), particularly due to the tapered posterior body that helps to prevent separation (3, 11, 12, 19), the potential for thrust production on the anterior body of a swimming fish has not been examined experimentally. Some previous researchers hypothesized that fish could benefit from this effect, with local thrust greatly reducing the impact of the net drag expected on a carangiform swimmer’s anterior body (20). Indeed, in computational models, one can see areas of negative pressure on the anterior body (21, 22), but this effect has never been studied systematically or in living fishes. We therefore used a recent set of tools (23, 24) to quantify the pressure and forces produced during swimming for two fish species that both have airfoil-shaped anterior bodies, bluegill sunfish (*Lepomis macrochirus* Rafinesque) and brook trout (*Salvelinus fontinalis* Mitchill), at high temporal and spatial resolution, the first such experimental test for negative pressure thrust production in living carangiform swimmers.

It is known that some fishes can produce negative pressures during swimming. Specifically, Gemmell et al. (25, 26) quantified the pressure distribution around larval lampreys and found that they produce negative pressures along the anterior parts of their bodies, resulting in thrust forces. In essence, larval lampreys suck themselves forward.

The negative pressures produced by larval lampreys are not due to airfoil-like mechanics. Instead, they are likely due to the high amplitude movements of their bodies (26), a pattern called “anguilliform” swimming, which is used primarily by a few eel-like elongate fish species (17, 27). Many anguilliform swimmers undulate a large fraction of their bodies at high amplitude, which is different from the pattern seen in many other fishes, which use the carangiform mode (17, 27). Moreover, larval lampreys use unusually high amplitudes when they swim, even compared to adult lampreys (28). It is not known whether negative pressure thrust is a quirk of their specific swimming mode, or whether such negative pressures can be produced by other fish species and swimming modes, particularly the carangiform mode, the most common swimming mode (27, 29).

We find that both bluegill sunfish and brook trout produce negative pressure thrust on their anterior bodies, but they do it using a very different mechanism from larval lampreys: the combination of their airfoil-shaped bodies and leading-edge suction. Our descriptions of pressure and force along the body also enable us to begin to tease apart how subtle differences in shape and movement affect swimming in a broader context. The carangiform swimming pattern belies the subtler but substantial variation in forms, movements, and ecological roles that exists within this mode (7, 20, 29, 30). For example, bluegill have a relatively deep trunk and shallow peduncle when viewed laterally, undulate only the posterior third of their bodies at a large amplitude (20), and are found in lakes, where they generally tend to hover or swim slowly (31, 32). In comparison, brook trout have a relatively shallower trunk and deeper peduncle, undulate slightly more of their body at large amplitude (20), and live in running water where they swim often and at high speeds (33, 34). These differences are sufficiently large that based on undulation amplitude alone, sometimes these fishes are considered examples of the two different carangiform subtypes – true carangiform (bluegill) and subcarangiform (trout) (30). These species differ in body shape and swimming movements; we identify subtle features of the swimming kinematics that lead to differences in their force production.

More broadly, our understanding of fish evolution and ecology is limited by the lack of comprehensive descriptions of swimming force production. Such descriptions, such as those presented here, will enable us to evaluate the strength of the relationships between body shape, movements, and swimming abilities. By helping to identify specific selection pressures underlying the diversity of modern fish forms, we can make predictions about the roles of different fishes within a given assemblage – species co-occurring in the same water body (7, 14, 18, 20). This understanding of the links between form and function in fishes can offer potential solutions for current underwater vehicle design challenges (35–37), such as producing animal-like vehicles less disruptive to aquatic life, enhancing swimming efficiency of biomimetic vehicles for longer-term deployments, or improving maneuvering capabilities for navigating environments with complex physical structure.

## Results

We measured fluid flow patterns in a horizontal plane around 5 bluegill sunfish (9.3-11.5 cm total length *L*) and 3 brook trout (10.0-11.0 cm total length) using standard digital particle image velocimetry (38). Individual fishes swam in a flow tunnel at 2.5 *L* s^-1^, which corresponded to Reynolds numbers (*Re = ρuL/μ*, where *ρ* is water density, *u* is flow velocity, *L* is fish body length, and *μ* is water’s dynamic viscosity) (17) of 20,000-30,000. Tailbeat frequencies were 4.9±0.5 Hz for bluegill and 4.7±1.0 Hz for trout, corresponding to Strouhal numbers (*St = fA/u*, where *f* is tailbeat frequency and *A* is peak-to-peak tailbeat amplitude) (22) of 0.156-0.404 and reduced frequencies *f*\* *=fL/u* (17) of 2.0±0.2 for bluegill and 2.1±0.4 for trout.

### The anterior body makes small movements

For both species, the amplitude (the distance from the center line to maximum excursion on one side or the other) was very small in the anterior body and increased in more posterior segments (Fig. 2A,B). For both species, in segments 1-3 (0-40% *L*), the amplitude was less than 2% *L*, and only increased to 3% *L* in segment 4 of trout and segment 5 of bluegill, before increasing to 6% *L* or more in the posterior-most segments (Fig. 2C,D). In comparison, the body’s maximum width was ∼13% *L* for both species (Fig. 2A,B). Likewise, the body angle made with the fish’s trajectory was less than 5° in segments 1-3 and increased over the posterior body to 30-40° (Fig. 2E,F).

**Fig. 2.**
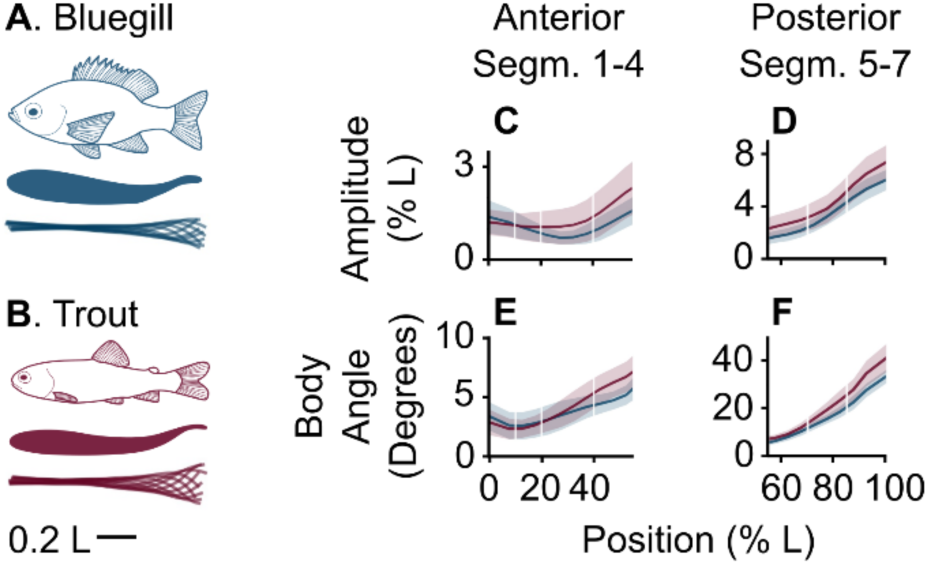
Bluegill and trout are carangiform swimmers. Overall midline kinematics for bluegill (A) and trout (B) swimming at 2.5 *L* s^-1^ indicate they are carangiform swimmers. Subtle differences between carangiform subtypes are visible through comparison of amplitude (C, D) and body angle (E, F) for anterior (C, E) and posterior (D, F) segments (Segm.), where segments are defined as in Fig. 3.

### The anterior body generates negative pressures

The body and tail motion swept fluid alongside the anterior body, like an airfoil, before accelerating the fluid alongside the posterior body and entraining it into vortices that were shed as the tail reached maximum excursion and changed direction (Movies S1, S2). This led to pressure fields (Fig. 3A,B, Movies S3, S4) with a region of strong positive pressure upstream of the snout, negative pressure along most of the anterior body, and oscillating positive and negative pressure gradients along the posterior body and caudal fin.

**Fig. 3.**
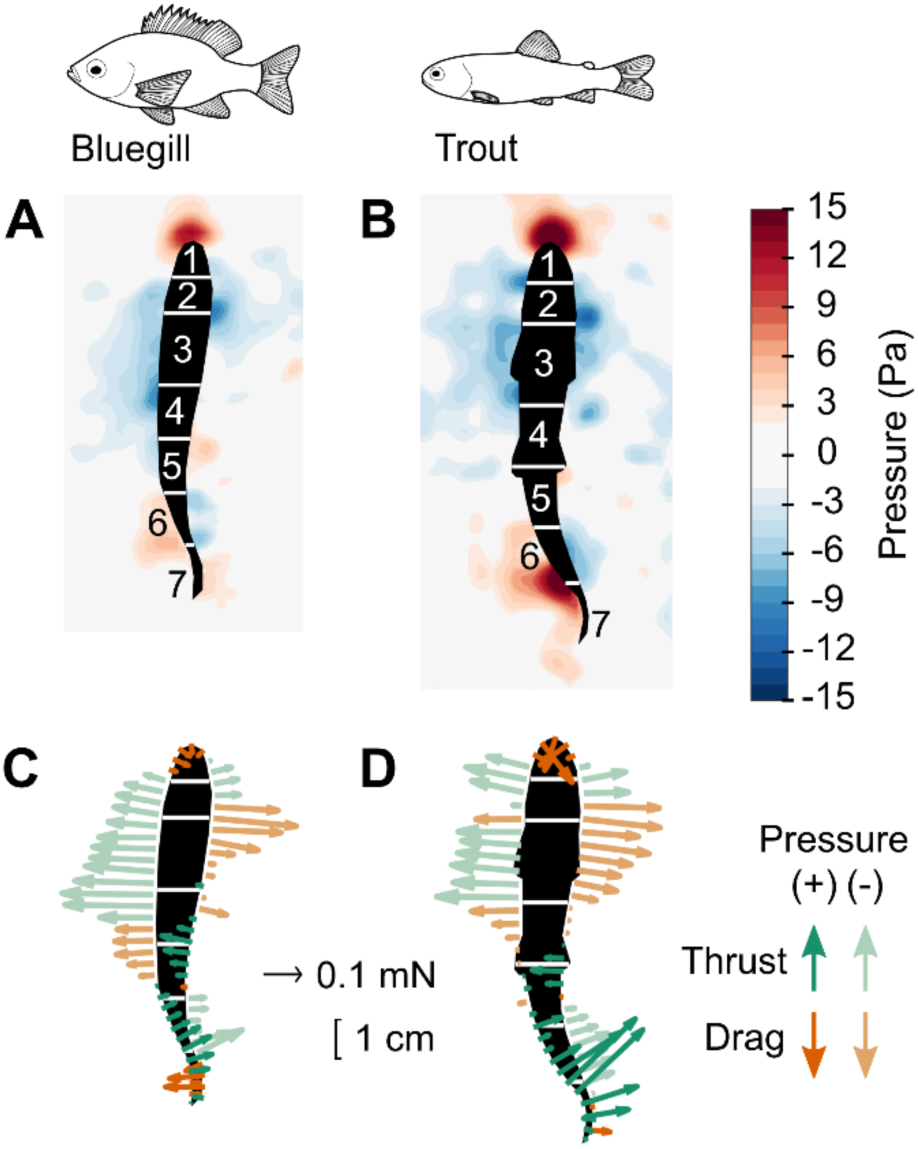
Undulatory swimming motions produce spatially and temporally complex patterns of pressure and forces. Panels show pressure fields (A, B) and estimated force vectors (C, D) for bluegill sunfish and trout swimming steadily at 2.5 *L* s^-1^. Numbering for body segments used throughout are given in (A, B), and the white lines indicate segment boundaries. For clarity only every third force vector is plotted in C and D.

To control for the difference in swimming speed among the fishes, we computed pressure coefficients: *C*_*P*_ = *P*/(0.5*ρu*^2^), where *P* is pressure. Fig. 4 shows the instantaneous pressure coefficients along one side of the body, along with the time-averaged value.

**Fig. 4.**
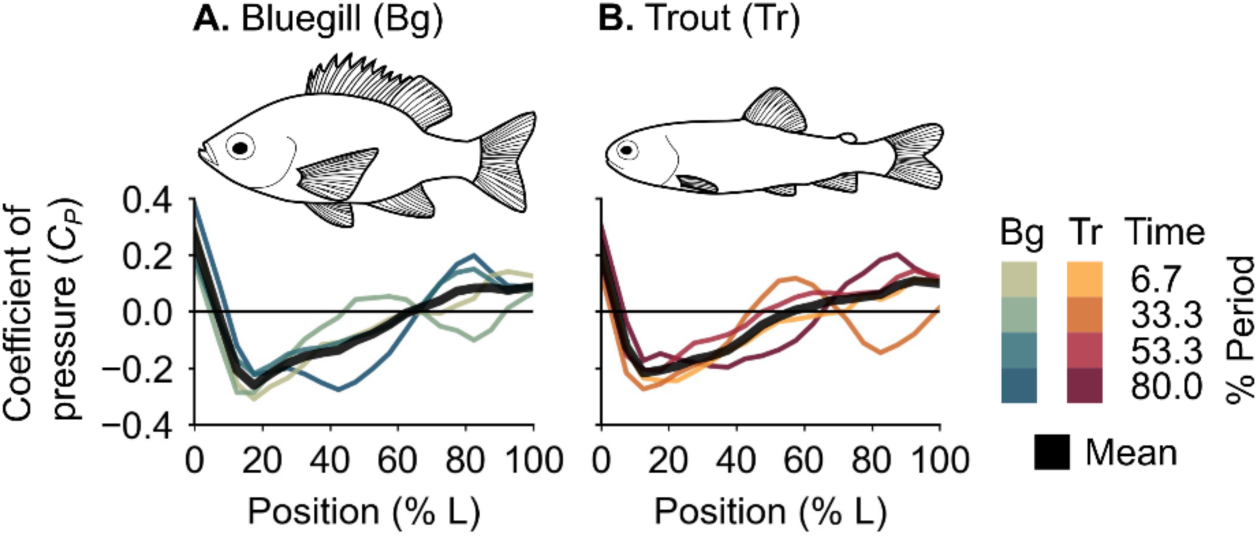
Profiles of pressure coefficient along the body vary over the tailbeat period. Colored traces show instantaneous profiles for bluegill (A) and trout (B) along the one side of the body, while thick black traces show the time-averaged mean.

The overall shapes of the pressure coefficient profiles had three important differences across the species (Fig. 4). First, the region of positive pressure on the snout was smaller in trout, resulting in negative pressure developing more anteriorly (see also Fig. 3A,B; Movies S3, S4). Second, bluegill often had larger magnitude negative pressure coefficients in in the midbody (10-55% *L*) than trout, but trout had larger positive and negative pressure coefficients in the posterior body (55-100% *L*). Finally, for both species, pressure shifted from negative in the midbody to positive near the tail, but for trout, this shift at times occurred more anteriorly (particularly at time t = 80.0% of the tailbeat cycle in Fig. 4).

The instantaneous pressure coefficients often differed greatly from the mean profiles (Fig. 4). Notably, the location where pressure coefficient changes sign from negative to positive shifted in the midbody region, and at times, a second area of negative pressure appeared on the posterior body (e.g., time t = 53.3% of the tailbeat cycle, Fig. 4).

### Negative pressure produces thrust on the anterior body

The shifting pressure gradients, combined with the body kinematics, led to complex spatial and temporal patterns of axial forces (Figs 3C,D, 5; Movies S5, S6). Both positive and negative pressure could produce thrust or drag, depending on the orientation of the body (see Fig. 1B). Thus, there were four types of forces: thrust due to positive pressure, thrust due to negative pressure, drag due to positive pressure, and drag due to negative pressure (Figs 3, 5). For bluegill, mean thrust forces were 1.3±0.5 mN. Trout produced a mean thrust of 1.5±0.4 mN. All values were on the same order of magnitude as previous estimates from wake analyses (39, 40).

**Fig. 5.**
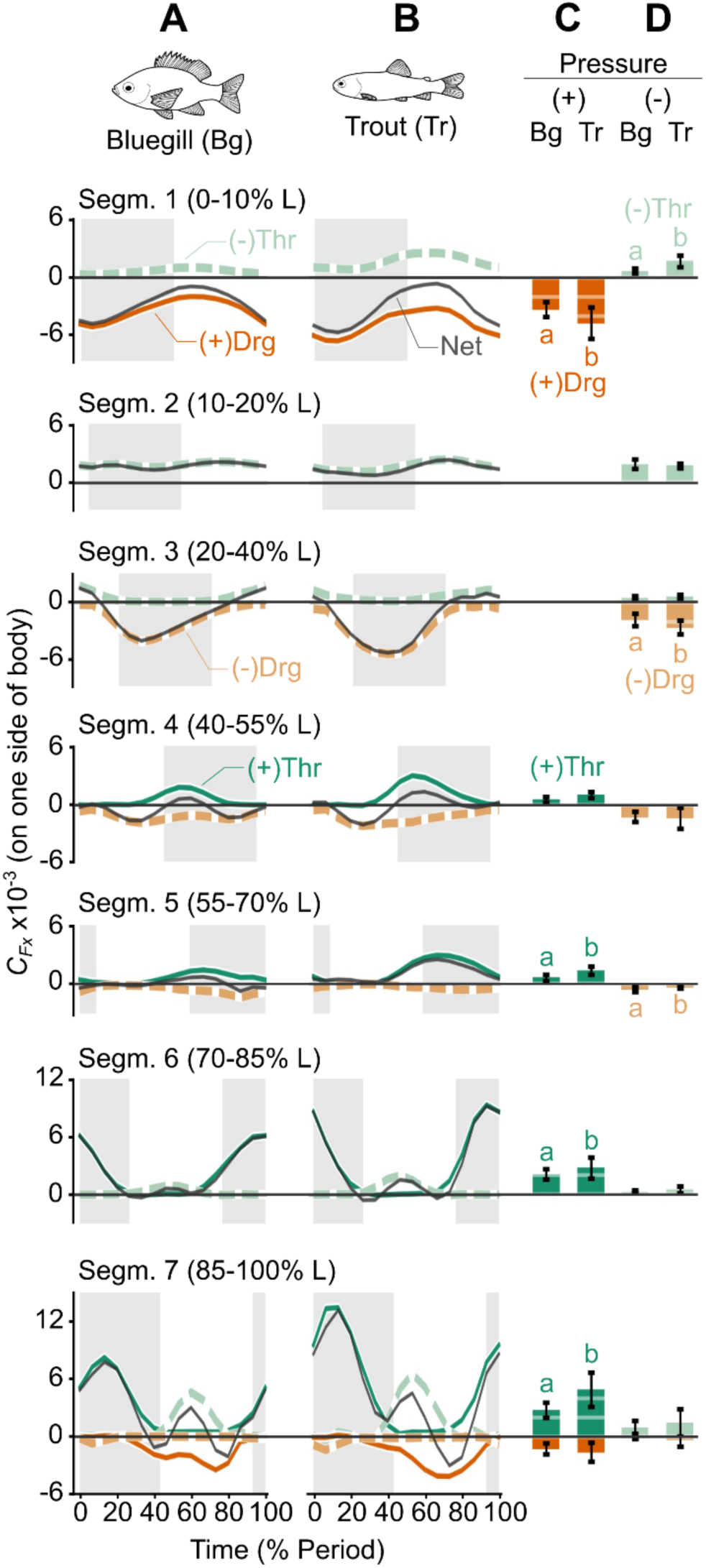
Thrust and drag arise from both positive and negative pressure in time- and space-dependent patterns. Panels A and B compare phase-resolved forces for bluegill (A) and trout for seven segments (Segm.) along one side of the body. Letters indicate where significant differences in force magnitude were detected across species (p < 0.05). The shaded region in the background indicates the times when the body segment moved from left to right, from peak amplitude to peak amplitude. Panels C and D compare mean thrust (Thr) and drag (Drg) forces arising from positive (C, (+)) or negative (D, (-)) pressure. When lines or bars are not shown, it means that both species’ mean force coefficients were effectively zero (*C*_*Fx*_ < 5% total *C*_*Fx*_ for that force type).

Fig. 5 shows the spatial and temporal patterns of these four forces in the two species, along with time-averaged values, on one side of the fishes’ bodies. Again, to control for the difference in body shape and swimming speed among species, forces were normalized to coefficients: *C*_*F*_ = *F*/(0.5*ρSu*^2^), where F is force and *S* is lateral surface area. Most of the mean coefficients for axial force subtypes were significantly different between bluegill and trout (Fig. 5C,D). Traces showing mean force coefficients summed across both sides of the body, as well as mean streamwise (total, rather than broken down by subtype) and lateral force coefficients on each body segment, are available in the Supporting Information (Figs S1-S3).

Spatially, the anterior body’s angle (Fig. 2E,F) combined with the negative pressures led to thrust on the anterior body and the tail, while positive pressures contributed to thrust only in posterior segments (Fig. 5). On the tip of the snout, positive pressure produced net drag (dark orange), but slightly more posteriorly, the pressure became negative, producing negative pressure thrust (light green). This shift occurred in segment 1 (0-10% *L*) for trout but in segment 2 (10-20% *L*) for bluegill, and in both species, negative pressures in segment 2 (10-20% *L*) produced thrust. Positive pressure thrust coefficients (dark green) occurred in segments 4-7 (40-100% *L*) and increased from anterior to posterior. Negative pressure thrust (light green) also occurred in the most posterior segments (segments 6-7, 70-100% *L*). Positive pressure drag (dark orange) was only present in segments 1 (0-10% *L*) and 7 (85-100% *L*). Negative pressure drag (light orange) was concentrated in the midbody (segments 3-5, 20-70% *L*).

### Trout produce positive pressure thrust more anteriorly than bluegill

The pattern of axial force coefficients along the body was different among species, depending on whether force was a thrust or drag force, and whether the force came from positive or negative pressure (linear mixed model ANOVAs: significant four-way interaction among species, force type, pressure type, and body segment; numerator DF = 6, denominator DF = 610, F-value = 4.1312, p = 0.0004). Where the two species had significantly different force coefficients, trout had larger magnitudes than bluegill, except for negative pressure drag in segment 5 (55-70% *L*, Fig. 5).

Fig. 6 compares within a species the different force types in three posterior segments that are functionally important. For bluegill, segment 4 (40-55% *L*) had significantly more drag than thrust (Fig. 6A), but in trout these two forces were equal (Fig. 6B). In segment 5 (55-70% *L*), the pattern shifted; trout produced more thrust than drag (Fig. 6B), but in bluegill they were equal (Fig. 6A). Thus, bluegill produced net drag in segment 4 and no net force in segment 5, while trout produced no net force in segment 4 and thrust in segment 5 (Figs 6, S2). Moreover, trout produced the same amount of lateral force as bluegill in segment 5 (Fig. S3). The kinematics of these segments were different in the two species: trout had higher amplitudes and higher angle to the horizontal (Fig. 2).

**Fig. 6.**
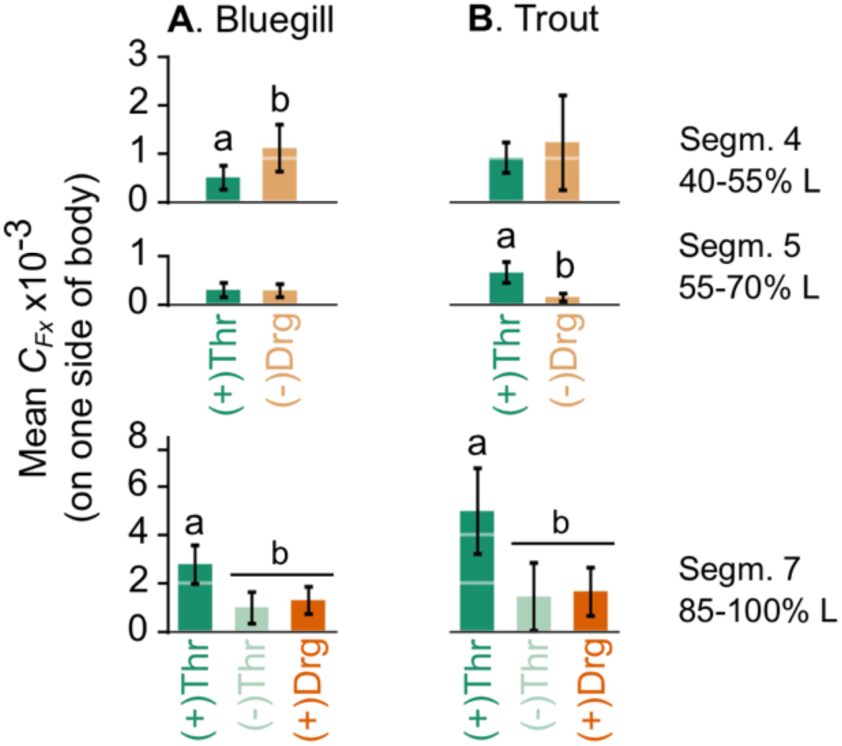
Bluegill and trout use their bodies differently to produce swimming forces. Comparison of thrust (Thr) and drag (Drg) forces from positive (+) and negative (-) pressure on one side of the body at posterior body segments (Segm.) for bluegill (A) and trout (B). Letters indicate significant differences between force types within a species (p < 0.05).

We approximate hydrodynamic Froude efficiency *η*, the ratio of useful power to total power (17), as *η* = ∑_*i*_(**F**_**T**,*i*_ · **v**_*i*_)/∑_*i*_| **F**_*i*_ · **v**_*i*_| where **F**_**T**,*i*_ is the thrust force vector, **F**_*i*_ is the total force vector, and **v**_*i*_ is the total velocity relative to the flow (including both side to side motion and the flow velocity) each on segment *i*. Based on this estimate, trout swim with an efficiency of 29.5±1.9%, compared to 26.6±1.0% in bluegill (mean±standard error; no significant difference across species; p = 0.142).

## Discussion

Many mechanical explanations of fish swimming emphasize that fishes push fluid behind them as they swim, creating areas of positive pressure on the body that push the fish forward as thrust forces (17, 25, 37). Thus, the recent discovery that larval lampreys rely substantially on negative pressure for thrust production (25) pointed to the underappreciated role of negative pressure in fish locomotion. Here, we present experimental data to show that the shape and oscillation of the airfoil-like body, common to many species of fishes, results in negative pressures that contribute significantly to thrust through a different mechanism than that used by larval lampreys. Using recent techniques for temporally- and spatially-resolved pressure and force measurements (23, 24), we find that, like in lampreys, negative pressure contributes significantly to swimming forces along a carangiform swimmer’s body (Fig. 5), producing 39% of the total thrust over the whole body. Unlike lampreys, however, most of the negative pressure thrust produced by carangiform swimmers arises not from high-amplitude swimming motions, but rather from the airfoil-like mechanics of the anterior body. Negative pressure acting on the anterior body produces 28% of total thrust. For comparison, the anterior body produces 36% of the total thrust when positive and negative pressure contributions are combined.

In addition, the high spatial and temporal resolution of our methods allows us to determine how small differences in kinematics among swimmers produced significant differences in forces (Figs 2, 6). Specifically, small differences in the body amplitude and angle of the posterior body, in combination with differences in lateral body depth profiles, allowed trout to produce higher thrust forces without increasing lateral forces and so may allow them to swim more effectively than bluegill. Thus, control of pressure gradients via both the airfoil-like shape of the anterior body and the kinematics of the posterior body are important for the effective development of swimming forces.

### Thrust on the posterior body comes from both positive and negative pressure

About two-thirds of the thrust comes from a familiar undulatory mechanism, as predicted by earlier studies (17, 20, 41, 42), relying on both positive and negative pressure in the posterior body. Time-averaged pressure profiles were previously measured by Dubois et al. (42) and theirs and ours both generally resembled the time-averaged pressure pattern on a pitching airfoil (Fig. 1A), especially on the anterior half of the body. Our profiles from the posterior body only look like theirs when averaged over a tailbeat cycle (Fig. 4). In instantaneous pressure profiles, pressure changes sign depending on location on the body and time within the tailbeat cycle (Fig. 4), resembling the distinct, alternating “pressure” (positive pressure) and “suction” (negative pressure) regions alongside the posterior body posited by Müller et al. (41) and found on the posterior bodies in computational models of carangiform swimmers (21, 22). This contrasts with the uniformly negative pressure on the posterior portion of a pitching airfoil (Fig. 1A). In particular, the caudal fin (segment 7, 85-100% *L*) experienced three forces: positive pressure thrust on the leading side of the lateral motion, negative pressure thrust on the trailing side, and positive pressure drag on the trailing side (Fig. 5). Together, these three forces produce a peak in thrust every time the caudal fin travels between peak excursions and near-zero forces as the caudal fin changes direction (Figs 5, S1). Dubois and colleagues (42–44) were not unaware of these effects; they noted that pressures fluctuated on some parts of the fish’s body in rhythm with the tailbeat, that there were negative pressures on the trailing side of the caudal fin, and that the caudal fin produces some drag, observations that all agree with ours.

We suggest that the actions of all three of these forces are necessary to create the shape of the characteristic double-peak pattern of thrust production over a tailbeat cycle (20, 29, 43–45). The positive pressure acting on the leading side of the caudal fin (segment 7, 85-100% *L*) is the primary source of thrust, leading to the magnitude of peak forces in the net force curves (Fig 5A,B, S1), since the magnitude of negative pressure thrust is equal to the magnitude of positive pressure drag (Fig. 6). This, again, agrees with computational models of carangiform swimming, where thrust was concentrated on the caudal region (21, 22). But, the timing of the peaks in positive pressure thrust on the leading side of the caudal fin and in negative pressure thrust on the trailing side differs (Fig. 5A,B, S1). And, the staggered timing of negative pressure thrust and positive pressure drag on the caudal fin – with the thrust acting first and quickly, and the drag acting second and slowly (Fig. 5A,B) – influences the timing of peak thrust and the shape of the net force curve on the caudal fin. This influence is visible when comparing across bluegill and trout; in trout, the negative pressure thrust peak occurs earlier, leading to net force curves with different shapes across species (Figs 5A,B, S1).

The implication here is that a fish’s control of pressure gradients around its caudal fin through adjustments to caudal fin shape or body kinematics may be vital for tuning thrust production on the posterior body. This agrees with Müller et al.’s (41, 46) hypothesis that fish can make small adjustments to their kinematics to control flow around the body and fine-tune their swimming performance, and further, this points to specific features – caudal fin shape and posterior body kinematics – that could have been influenced by selection on swimming abilities over the course of fish evolution.

It is important to note that the patterns of force production we describe here only reflect steady swimming. Presumably, timing, magnitude, and location of forces, in addition to the relative role of positive and negative pressure, could all change during accelerations. For example, many carangiform swimmers, including bluegill, have larger head and tail oscillation amplitudes and larger tailbeat frequencies during accelerations (39, 47), leading to larger added masses and larger total forces (39). Interestingly, in bluegill (39) but not trout (47), these increases occur without substantially redirecting the net thrust forces relative to steady swimming, suggesting that there are differences in the force production mechanics among species and across behaviors like steady swimming and accelerations.

### Trout may produce swimming forces more effectively than bluegill

From their lifestyles, we might hypothesize that bluegill, which generally hover or swim slowly in still water or slowly-flowing streams (20, 31, 32), do not produce thrust as effectively as trout, which spend much of their lives swimming (20, 33, 34), even though both swim in a similar way. If this hypothesis is correct, then what aspects of kinematics or body morphology in trout lead to more effective swimming? Answering questions like these, both within and across swimming modes, would allow us to evaluate the strength of relationships between swimming abilities, morphology, and kinematics, and further, identify specific selection pressures that may have led to modern fish forms. Our pressure measurements allow us to approximate hydrodynamic Froude efficiency, the ratio of useful and total power. We find that the Froude efficiency is 2.9% higher in trout than bluegill. Earlier predictions likewise suggest that trout may have higher Froude efficiencies than bluegill (20). The difference we observe in efficiency, while not significant (p = 0.142), may point toward functional differences in thrust production between trout and bluegill. Froude efficiency is only a mechanical efficiency, and does not account for potential differences in metabolic rates (1), but even such small differences in efficiency could lead to significant energy savings over the long bouts of continuous swimming typical of a trout’s lifestyle (20, 33, 34).

Indeed, we hypothesize that the subtle differences in kinematics and body shape among the species are functionally meaningful. The midbody (segments 4-5, 40-70% *L*), where forces transition from drag to thrust, is the most functionally relevant. In bluegill, the transition from drag to thrust occurred on body segment 5 (55-70% *L*), where the net force coefficient was near zero (Figs 6A, S2). In contrast, in trout, this transition occurred more anteriorly in segment 4 (40-55% *L*), with segment 5 (55-70% *L*) clearly producing thrust (Figs 6B, S2).

These differences seem to reflect kinematic differences among the species: trout are sometimes classified as “subcarangiform” swimmers, which have higher amplitude undulations more anteriorly on their body than “true-carangiform” swimmers like bluegill (Fig. 2A-C) (20). First, the more anterior transition to undulatory motion in trout means that the development of thrust-producing positive pressure gradients occurs more anteriorly, too (Fig. 4, time t = 53.3 and 80% of the tailbeat cycle). Second, in trout, these more posterior segments make a larger angle to the swimming trajectory (Fig. 2E,F), directing the forces more toward thrust than lateral forces. Indeed, the ratio of axial to lateral force coefficients is much larger in this segment in trout than in bluegill (0.33 in trout and 0.06 in bluegill) (Figs 5, S2, S3).

As a whole, our results suggest that trout are producing swimming force more effectively than bluegill. This is because they produce higher thrust forces than bluegill and use more of their body to produce thrust. But, although trout are undulating at larger amplitudes, the lateral forces they produce are no different from or are less than (segment 4, 40-55% *L*) those of bluegill (Fig. S3). Since lateral forces are wasted effort (part of the denominator in Froude efficiency), these larger body undulations do not appear to be incurring additional costs for the trout. We suggest that this is due to trout’s shallower body depth profile. While a full analysis of morphology, lateral forces, and swimming efficiency is beyond the scope of this study, these findings suggest that an examination of subtle differences across carangiform swimmers is a promising direction for future work linking form and function in fishes.

### The anterior body produces thrust due to airfoil-like mechanics

Despite the differences in force distribution in posterior segments, the overall pattern of pressure and forces in the anterior body is quite similar across bluegill and trout and much like that over an airfoil. The reduced frequency of oscillation is fairly large (∼2 for both species), suggesting that oscillatory mechanics might be more important than airfoil-like mechanics. However, we find that the pressure distribution on the anterior body is very similar to an airfoil at a constant angle of attack (reduced frequency of 0) (48) or a pitching airfoil at a much lower reduced frequency (0.2 in Fig. 1A) (13).

For both fishes, the cross-sectional shape of the anterior body is close to that of a NACA airfoil (Figs 1A, 2A,B) (2, 3), leading it to develop negative pressure along most of its length (Figs 3, 4), like an airfoil (Fig. 1A) (4, 11). In both fish species, as in airfoils with an angle of attack to the flow (Fig. 1A,C) (11, 12, 14–16), the region of positive pressure is not directly on the tip of the snout (Fig. 4; Movies S3, S4). Instead, it oscillates to either side (Figs 1A, 4; Movies S3, S4), and the rest of the anterior body (segments 2-3, 10-40% *L*) solely experiences negative pressure (Figs 3, 4). This process is similar to leading-edge suction mechanics on airfoils at moderate angles of attack (Fig. 1C) (11, 12, 14–16). Our observations of negative pressure also match measurements by Dubois et al. (42, 44), who implanted pressure cannulae under the skin of bluefish and found that negative pressure dominated much of the bluefish’s length, leading to mean pressure profiles shaped similarly to those in Fig. 4 and suction-based thrust forces on the anterior body. Likewise, we find that negative pressure – arising from the airfoil-like shape, and placed far forward on the anterior body due to leading-edge suction mechanics – is positioned alongside forward-facing body surfaces and leads to small but significant, continuous thrust in segment 2 (10-20% *L*) (Figs 3, 5A,B).

Similar pressure distributions have also been found in 3D computational fluid dynamics models of carangiform swimmers. Borazjani and Sotiropoulos (21) and Liu et al. (22) both documented negative pressure regions along the anterior bodies in simulations of mackerel and crevalle jack, respectively, but they did not highlight the role of these negative pressures in thrust production. Even so, the presence of these mechanics across five phylogenetically-distant species points to the ubiquity of airfoil-like thrust generation among carangiform swimmers.

This thrust production mechanism means that the anterior body produces less drag than it might otherwise, but it is still net drag producing. Dubois et al. (42–44), Anderson et al. (19), Borazjani and Sotiropoulos (21), and Liu et al. (22) all find that the anterior body produces net drag forces. Our work does not contradict these findings; indeed, we find that, on the anterior body, the magnitude of the negative pressure thrust forces is smaller than the sum of drag forces (positive pressure drag on the tip of the snout, segment 1, 0-10% L, and negative pressure drag on the midbody, segment 3, 20-40% L) (Figs 5, S2). However, the negative pressure thrust on the anterior body (segments 1-4, 0-55% *L*) balances out a large fraction (45%) of this drag, causing the anterior body to produce much less net drag. Thus, we point to a more nuanced role for the anterior body during carangiform locomotion, as anterior-body thrust forces make up a substantial portion of the total thrust.

These thrust forces arise from very small movements of the anterior body (Fig. 2A-D), and likely require little muscle activity. At low speeds, like in this study, trout do not activate red muscle anterior to 50% *L* (49), and neither do largemouth bass, a species closely related to bluegill sunfish (50). Thus, the small, mostly passive movement in the anterior body may allow this airfoil-like thrust production to be highly efficient.

This airfoil-like mechanism is different from the mechanism Gemmell et al. (25, 26) identified in larval lampreys, which also produce thrust due to negative pressure (Fig. 7). Larval lampreys swim in the anguilliform mode, which is characterized by large-amplitude undulations in the anterior regions of the body (Fig. 7) (17, 25, 27). Even among anguilliform swimmers, the larval lampreys studied by Gemmell et al. (25, 26) have particularly large anterior body movements (28). These undulations rotate the body surface, which accelerate the adjacent fluid, strengthen the fluid’s vorticity, and generate large regions of negative pressure (Fig. 7A) (26, 51, 52). This negative pressure leads to suction-like thrust forces, which act continuously along much of the length of the body (Fig. 7A) (25, 26, 51, 52). In contrast, bluegill and trout, which are carangiform swimmers, produce negative pressure locally on their anterior bodies due to their cross-sectional shape and small motions (Fig. 7B).

**Fig. 7.**
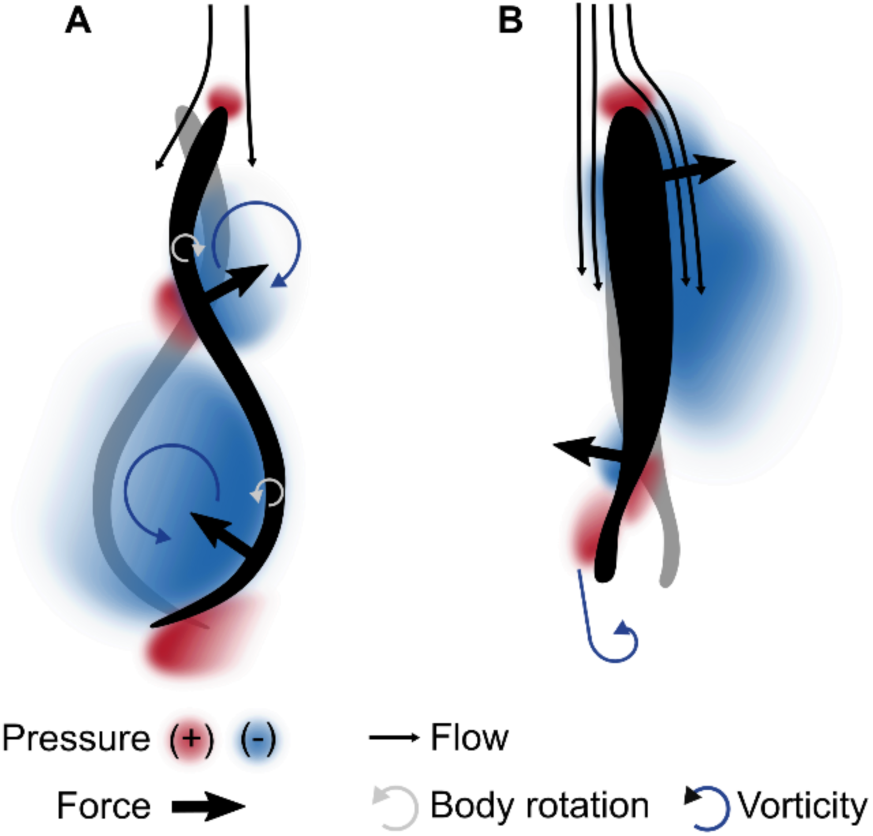
Anterior body thrust for anguilliform and carangiform swimmers is based on different mechanics. Gray and black silhouettes show the motion of the body, and color indicates pressure. Anguilliform swimmers (A) produce negative pressure thrust along the whole body using an undulatory pump mechanism, in which high amplitude body movements suck fluid along the body. Anguilliform kinematics adapted from (57). In contrast, carangiform swimmers (B) produce thrust on the anterior body through airfoil-like mechanics. For clarity, only negative pressure thrust forces are shown.

A growing body of work points to how different swimming modes and body shapes most likely confer different functional advantages (5–10). Detailed comparisons of force production by specific parts of the fish body like those performed here will allow us to finally test these hypotheses, and ultimately, arrive at a more complete understanding fish evolution and ecology. For example, we have long hypothesized that streamlined bodies like those of a tuna enable the fast, efficient swimming required of Pacific migrations. Here, we show that this body streamlining may contribute to the efficiency of thrust production. These fishes not only produce low drag but can also take advantage of the airfoil-like cross-section of their body and recoiling movements to produce thrust. Because the streamlined body cross-section and small anterior-body oscillations are very common in fishes, we suggest that this mechanism of producing thrust might be a general feature of swimming in many fish species.

## Materials and Methods

Full details of the methods can be found as SI Materials and Methods.

### Experimental procedures

Individual bluegill and brook trout swam at 2.5 *L* s^-1^ in a recirculating flume seeded with near-neutrally buoyant particles illuminated by horizontal laser light sheets from two sides. Fishes were filmed using two high-speed cameras (Photron Fastcam Mini AX50, 1024 x 1024 pixel resolution, 20 μm pixel size), which captured synchronized ventral and lateral view footage at 1000 and 100 frames per second, respectively. Only sequences where the fish used steady, body-caudal fin swimming motions for at least 1.5 tailbeats cycles within the light sheet were processed. Video of 3 replicate swimming trials was collected for each individual. Experiments were approved by the Harvard University Institutional Animal Care and Use Committee under protocol 20-03 (GVL).

### Data processing

Water velocity was calculated using particle image velocimetry (PIV) in DaVis 8.2.2 (LaVision GmbH, Goettingen, GER), with interrogation window sizes 32 x 32 pix and 16 x 16 pix, 50% overlap, and two passes at each window size (38).

Following our previously-validated protocol (24), ventral outlines of the fish were manually digitized in ImageJ (NIH, Bethesda, MD, USA). Midlines were extracted automatically from these outlines using a custom Matlab 2015b (Mathworks, Inc., Natick, MA, USA) script. Midline kinematics (e.g., tailbeat period, frequency, lateral amplitude, and body angle) were calculated using a custom script in Python (version 2.7.11, Python Software Foundation; https://www.python.org) following Videler (29). We use the mathematical amplitude, the distance between the center line and maximum lateral excursion, which is half of the peak-to-peak lateral excursion, often referred to as “amplitude” in older works (1, 53). To facilitate comparisons across different parts of the fishes’ bodies, fishes were divided into seven body segments which grouped together portions of the body with similar kinematics, body shape, and pressure gradients. Pressure and forces calculated below were averaged within segments.

Pressure distributions were calculated following Dabiri et al. (23) in Matlab using velocity data and outlines of the fishes’ bodies. We estimated forces using the procedure detailed in Lucas et al. (24). In brief, force magnitude was calculated as the product of pressure and surface area at a point in a calculation boundary drawn around the fish, where the area was the product of the distance between points in the horizontal plane and the fish’s body depth at those points. Force vectors were directed inward or outward based on the sign of the surrounding pressure. Our previous validations (24) indicate that, for fish-like swimmers, pressure effects dominate shear effects (e.g., skin friction), and that this 2D approach is robust to the out-of-plane flows around a fish, e.g., Liu et al. (22), allowing for accurate estimation of forces through these procedures.

### Statistics

Linear mixed effects models were developed following the standard practice outlined by Zuur et al. (54). For axial forces (*CFx*), two models were developed. The first compared the mean magnitudes of axial force subtypes and included four fixed effects, each with multiple levels: force type (thrust, drag), pressure type (positive, negative), species (bluegill, trout), and segment (1-7), and all interactions between these effects. The second model examined the means of total axial forces. Both this model and the model for mean lateral forces (*CFy*) included two fixed effects: species and segment, and their interaction. The model for efficiency included one fixed effect: species. In all models, individual was included as a random effect, and in force models, variance specifications accounted for heterogeneity (54, 55). ANOVA tests and post-hoc pairwise comparisons were conducted to determine which effects significantly affected force coefficients. A false discovery rate correction was applied to all post-hoc results (56).

All statistics were performed in R (version 3.5.1, R Foundation for Statistical Computing, Vienna, Austria; https://www.r-project.org/) using the nlme package (version 3.1-137, https://CRAN.R-project.org/package=nlme), and marginal means were estimated for pairwise comparisons using the emmeans package (version 1.2.3, https://CRAN.R-project.org/package=emmeans).

## Acknowledgements

Thanks to Mariel Rosic for assistance with data collection; members of the Lauder Lab for fish care; John Dabiri, Brad Gemmell, and Gary Lucas for technical assistance; Tyler Wise for sample data used to develop code; Steve Worthington and Kara Feilich for statistical advice; Paul Webb, Sean Colin, Elena Kramer, Pete Girguis, and members of the Girguis and Tytell labs for helpful discussions; Karthik Menon and Rajat Mittal for airfoil surface pressure data; and the anonymous reviewers whose feedback greatly improved the manuscript. This work was supported by a National Science Foundation Graduate Research Fellowship under grant DGE-1745303 to KNL and by NSF IOS 1652582 to EDT, and by the Office of Naval Research Multi-University Research Initiative Grant N000141410533 monitored by Dr. Bob Brizzolara to GVL. The funders had no role in study design, data collection and analysis, decision to publish, or preparation of the manuscript. No competing interests declared.

