## Supplemental Information for "The fish body functions as an airfoil: surface pressures generate thrust during carangiform locomotion"

### SI Materials and Methods

##### Fishes

Bluegill were captured by beach seine from White Pond, Concord, MA, USA, and were housed individually in 38 L tanks kept at 20°C. Bluegill were fed a combination of live worms and pellets. Brook trout were purchased from Blue Stream Aquaculture, West Barnstable, MA, USA, and were maintained in a 1500 L recirculating tank kept at 16°C. Trout were fed 3.5 mm high protein pellets daily (Keystone Hatcheries, Richmond, IL, USA). Fishes were kept on a 12-hour light: 12-hour dark photoperiod. All fish care and experimental protocols were approved by the Harvard University Institutional Animal Care and Use Committee under protocol 20-03 (G. Lauder).

##### Experimental setup and video collection

Experiments were performed in a 600 L recirculating flume with a 28 x 28 x 80 cm working area. Fish were blocked from drifting downstream outside the test area by a baffle and were encouraged to hold position in the center of the tank by the presence of vertical black strips of plastic clamped to the walls of the tank. Intake and outflow pipes from a chiller unit were positioned downstream of the test section. Water was chilled to 16°C during trout testing but was maintained at room temperature (20°C) for bluegill. Flow rates in the flume were controlled using a custom LabVIEW program (National Instruments Corp., Austin, TX, USA).

The flume was seeded with near-neutrally buoyant (density 1060 kg $m^{-3}$) VESTOSINT 1164 white nylon 12 particles with an average diameter of 50 µm (Degussa Corporation,

Piscataway, NJ, USA; now Evonik Industries AG, Essen, GER), and flow was pulsed to high speeds periodically throughout testing to resuspend particles. Tracer particles were illuminated by two light sheets, one entering from either side of the tank and aligned to within 1 mm of each other, so as to eliminate shadows around the fish's body. Light sheets were produced by spreading the beams from two continuous wave 532 nm solid-state lasers (MGL-N-532A, OptoEngine, Midvale, UT, USA).

Before transfer to the test tank, each fish was measured for total length and then transferred to a small, rectangular container and photographed in lateral view with a scale bar. In the test tank, fish were given an acclimation window of 20 minutes to 12 hours prior to data collection, depending on the species, individual stress responses to handling, and response to the laser lights.

Individual swimming fish were filmed from two perspectives using two Photron Fastcam Mini AX50 (1024 x 1024 pixel resolution, 20 μm pixel size) high-speed video cameras. The first camera captured ventral view footage of the fish and surrounding flow off of a 45° mirror below the tank at 1000 frames per second (fps), and it was positioned to capture flow in the central region of the tank away from the walls, leading to images capturing a 26 x 26 cm space. The second camera was positioned adjacent to one of the lasers and aimed diagonally upstream. This camera filmed at 100 fps in sync with the ventral view camera and provided nearly-lateral view images of the fish, which were used to confirm that the fish was vertically centered in the laser light sheet during swimming trials selected for further processing. To ensure visibility of the fish, a red-pass filter (Schott Color Glass Filter, CG-OG-530-2.00-2.5, CVI Laser Corp., Albuquerque, NM, USA) was used on the lateral-view camera to reduce glare from the light sheet. Cameras ran continuously, and an end trigger was used to keep the last 12 seconds of video in memory for review and saving when the fish swam in the light sheet.

The criteria we used to select videos for processing are as follows. First, we required sequences where the fish used steady, body-caudal fin swimming motions for 1.5 tailbeat cycles. Although data would only be extracted from the duration of 1 tailbeat cycle during analysis, the extra 0.5 cycles were included to allow for a time buffer at the start and end of the sequence. We defined steady, body-caudal fin motion as the fish staying within 5 mm of its starting position (with one exception, where a 7 mm drift was permitted in order to maintain balanced experimental design) and not using pectoral fin beats. For each individual, footage of 3 replicate swimming trials was collected. Both fishes swam at 2.5 body lengths per second (L $s^{-1}$). Only sequences where the whole body remained in the light sheet for the duration of a tailbeat were selected for further processing.

Calibration images were produced each time the cameras were moved by placing a plate with crosses marked on a 1 cm square grid in the test section at the depth of the laser sheet and taking photographs with the camera positioned beneath the tank.

**Particle image velocimetry**

Particle image velocimetry (PIV) analysis was conducted in DaVis 8.2.2 (LaVision GmbH, Goettingen, GER) (1–3). Cross-correlation was conducted with decreasing interrogation window sizes (32 x 32 and 16 x 16) and 50% overlap. Two passes were made at each window size. During postprocessing, vectors were deleted if their correlation value was <0.8, and the empty spaces were filled by interpolation. This resulted in a 128 x 128 grid of velocity vectors. Due to the difficulty in automatically tracking fish fins, which appeared translucent and often in poor contrast to the background, fish were not masked during vector calculation, so vectors were calculated over the whole image, including inside of the fish's body. These internal vectors did not represent real flows. Smoothing regimes such as those used by Lucas et al. (4) consider the average flow among a vector's nearest neighbors (3). Because the fishes were not masked during processing, no smoothing regime was applied to prevent erroneous vectors calculated inside the fish's body from influencing real flow data.

**Digitization for kinematics, body depth, and lateral area**

Although ventral view videos were collected at 1000 fps for PIV analysis, our previous validations (4) indicated that major trends in forces on a swimming body are captured when pressure and force are calculated from velocity fields at 100 fps. For this reason, ventral outlines of the fish were manually digitized in ImageJ (NIH, Bethesda, MD, USA) in every tenth frame of ventral view video. In each outline, the first point was always placed on the anterior-most tip of the fish, and digitization thereafter proceeded clockwise, leading to outlines of 20-30 points. In addition to the kinematics tracking described below, ventral view outlines were also used in the calculation of pressure and forces.

Fish midlines were extracted automatically in a custom Matlab 2015b (Mathworks, Inc., Natick, MA, USA) script from manually-digitized ventral outlines. To do this, 20 points were initialized inside the fish outline, evenly spaced on the x-axis between the manually-digitized anterior-most point, and an automatically identified tail tip. The tail tip was extracted based on curvature of the outline near points with the maximum x-coordinates, as fish always swam toward the left side of the screen. This initial straight line of 20 points was adjusted into a midline using a custom snake algorithm (5), which iteratively moved the 18 internal points in small steps in a direction chosen based on the weight of three tendencies: a tendency to move away from the edges of the fish outline, a tendency to stay near other points, and a tendency not to make a sharp angle with neighboring points. An arc-length interpolation was then used to generate a midline with 100 equally-spaced points.

Midlines kinematics were then calculated using a custom script in Python (version 2.7.11, Python Software Foundation; https://www.python.org). To do this, the midline was first smoothed using a quintic least-squares spline. Tailbeat period and frequency were then extracted from midline motion by tracking inflection points (positions of zero curvature) in the kinematic waveform traveling along the posterior half of the body following the methods described in Videler (6). Lateral amplitude of the kinematic waveform was calculated as the mean of peak lateral excursion of the midline, and the body angle – the angle the body made with the fish's trajectory – was calculated as the tangential angle at each point on the midline.

In addition to the ventral outlines, a lateral view outline of each fish was manually digitized in ImageJ from the lateral view photographs taken before fishes were transferred into the test tank. The lateral area of a fish's body was measured as the area enclosed by its manually-digitized lateral outline. Using the same initialization and snake algorithm process applied to make ventral view midlines above, lateral view midlines were generated. Body depth could then be automatically measured along lines drawn perpendicularly to the lateral view midline. This was accomplished by finding the distance between the points where the perpendicular lines intersected the dorsal and ventral outlines of the fish. On the caudal fin lobes, the body depth was the sum of distances between the dorsal and ventral outlines on each lobe. A body depth was calculated for each ventral view midline point, using the distance-along-body measure to relate the ventral and lateral view images.

**Pressure and force calculation**

Ultimately, velocity fields generated from PIV were used to calculate pressure distributions in the water around the fish's body using the Dabiri et al. (7) algorithm. The Dabiri et al. (7) algorithm has been validated against computational simulations of flow around a square cylinder and an anguilliform swimmer and in both cases, captures the major pressure gradients around these bodies. These pressure fields were then used to calculate forces acting on the fish's body following our previously-described protocols (4). Our earlier work (4) also details the validations of these pressure-based force calculation methods and shows that for fish-like swimming, the effects of shear forces as small enough that a pressure-based calculation provides an accurate estimate of swimming forces. Further, 3D bodies in flow will create flows that are inherently 3D, as illustrated for fish in Liu et al. (8). These flows include tip vortices and may not be captured with 2D quantification techniques for bodies with the aspect ratio of a fish (9, 10). Our validations also examine the influence of these effects for fish-like swimmers, and we find that this 2D approach is able to accurately reproduce the shape, timing, and magnitude of force-vs-time curves of fish-like swimmers (4).

The following paragraphs describe the pressure and force calculation process in more detail. Prior to these calculations, two more pieces of information were needed: specifications of where the fish's body was in the images and of where in space pressures should be extracted for force calculation.

First, the manually-digitized ventral outlines were used to create masks for use with the Dabiri et al. (7) pressure field calculator. These masks would blank out velocity vectors inside the fish's body and indicated to the algorithm the presence of a solid body (4, 7). To ensure that velocity vectors were enclosed in the narrow rostrum and caudal regions so that pressures would not be calculated through the fish's body, digitized outlines were adjusted – extending the rostrum by 0.5 mm and widening the caudal fin by 1.5 mm (0.75 mm on either side) (4) in Matlab 2015b.

Following the processing sequence detailed in Lucas et al. (4), we then generated boundaries which specified where around the fish's body we were interested in pressure magnitudes. These boundaries were set in Matlab 2015b as 198-point loops encircling the fish within 2.5 boundary-layer-widths of its surface, where pressure was defined and forces could be calculated (4, 7). For each frame of video that would be processed, the boundary was generated in three steps. First, pelvic fins were removed from the body outline, as well as pectoral fins for trout (bluegill pectoral fins were held alongside the body). Paired fin outlines were required in the masks, as no flow information was available where these fins blocked the view of the laser light sheet, but for this same reason, forces could not be calculated directly on the portions of the body blocked from view by these fins. Then, an arc-length interpolation converted the remaining manually-digitized outline into an outline with 198 evenly-spaced points. Finally, this 198-point outline was expanded outward from the fish using another custom snake algorithm (5), which pushed the outline away from the fish while keeping boundary points relatively evenly spaced and the outline smooth.

We chose to calculate pressure in an 18 x 18 cm domain surrounding the fish, based on the following convergence analysis. Dabiri et al. (7) described in their supplemental material the need for care in choosing a domain size (length and width of the velocity field) because their algorithm calculates pressure along integration paths through the velocity field from the outer edge toward the center. Thus, domain size must compromise between keeping integration paths short to avoid accumulating error during pressure calculations and keeping the domain large enough so that the assumption that the pressure is zero on the edges of the domain was still valid. Following the protocol suggested by Dabiri et al. (7) to ensure the velocity field here met both of these criteria, a small sample of velocity fields were cropped to several dimensions (26, 22, 20, 18, 16, and 14 cm square fields and a rectangular 8 x 14 cm field cropped close to the fish's body). Then, the masks and these cropped velocity vector fields were then loaded into the Dabiri et al. (7) pressure algorithm to generate pressure fields. Pressure magnitudes on the calculation boundary and were plotted versus calculation boundary point to visualize fluctuations in the calculated pressure values induced by the changes in domain size. The 16, 18, and 20 cm square domains converged to similar calculated pressures around fish bodies – indicating minimal error induced by too large or too small domains – and so velocity fields were cropped to 18 x 18 cm windows centered around the fish during final pressure and force calculation.

After calculating pressure in this domain, we then estimated forces on the body. Force magnitudes were calculated at each point as the product of pressure and an area term, following the equations and procedure described in Lucas et al. (4). Pressures at force calculation boundary points located inside of paired fin outlines were estimated by linearly interpolating between pressures on either side of the fins. The area term for each calculation boundary point was the area of a rectangle whose width was the distance between calculation boundary points, and whose length was the depth of the fish's body at the calculation boundary point. Body depth for a calculation boundary point was assumed to be the body depth at the nearest ventral view midline point. Pressure-based forces always act perpendicularly to a surface, so force vectors were always parallel with the normal vector at the corresponding calculation boundary points. Force vectors were directed inward or outward based on the sign of the surrounding pressure – positive pressure pushes on the body surface and directs force inward, while negative pressure pulls and directs force outward.

To enable comparison across swimming speeds and species, pressure and force were both normalized to non-dimensional pressure and force coefficients ($C_p$ and $C_F$) as

$$C_p = \frac{p}{\frac{1}{2}\rho u^2} \quad (1)$$

$$C_F = \frac{F}{\frac{1}{2}\rho A u^2} \quad (2)$$

where $C_p$ is pressure coefficient, $p$ is pressure (Pa), $C_F$ is force coefficient, $F$ is force (N), $\rho$ is the density of fresh water (1000 kg $m^{-3}$), $u$ is the swimming speed (m $s^{-1}$), and $A$ is the lateral area of the fish's body ($m^2$).

Because fish entered the light sheet and swam continuously starting at different points in a tailbeat cycle, all data were synchronized for comparisons based on the movement of the tip of the caudal fin. All time-series were reordered so that at time t = 0 s the tail tip was positioned at its maximum excursion to the right side of the body (as viewed in the videos). To allow for averaging across trials, all data were downsampled to 15 time points evenly distributed across the tail beat period. The shortest tailbeat period observed was 0.15 s. In addition, the right and left sides of the body experienced the same pressures and forces, but mirrored and at a lag of half of tailbeat cycle. Thus, data from the left side was mirrored and synchronized with data from the right side, leading to 2 time-series from every replicate tailbeat cycle which were averaged together. Reported pressure and force data therefore reflect what happens on one side of the body, unless otherwise indicated.

To facilitate comparisons across different parts of the fishes' bodies, fishes were divided into seven body segments. These segments were defined so as to keep lengths of segments as even as possible while grouping together portions of the body with similar kinematics, body shape, and pressure gradients.

All kinematic, mean pressure, and net force information were aggregated and synchronized for data reporting and statistical tests using a custom Python script.

##### Hydrodynamic efficiency

We approximate hydrodynamic Froude efficiency $\eta$, the ratio of useful power to total power (11), as $\eta = \sum_i (\mathbf{F}_{\mathbf{T},i} \cdot \mathbf{v}_i) / \sum_i |\mathbf{F}_i \cdot \mathbf{v}_i|$, where $\mathbf{F}_{\mathbf{T},i}$ is the thrust force vector, $\mathbf{F}_i$ is the total force vector, and $\mathbf{v}_i$ is the total velocity relative to the flow (including both side to side motion and the flow velocity) each on segment $i$.

##### Statistics

Statistical models were developed to quantify how pressure and force differs along the body among the two species. This led to the use of linear mixed effects models relating mean magnitude of axial force coefficient ($C_{Fx}$) subtypes to four effects, each with multiple levels: force type (thrust, drag), pressure type (positive, negative), species (bluegill, trout), and segment (1-7), and all interactions between these effects. Individual was included as a random effect to account for natural variation between individual fishes (12, 13). Because an examination of the residuals indicated that $C_{Fx}$ data were heterogeneous, weights were applied to the model to allow for unequal variances among the grouping effects. Because all effects were categorical variables, unequal variance structures would allow for variance to differ between levels of one or more effects (12, 14). Appropriate variance structures were selected for each model by examining the variance and residuals at each level of each effect and constructing possible variance structures that described the unequal variance observed. The lowest Akaike Information Criterion score was used to select between models (12). After model selection, residuals were reexamined to verify that heterogeneity was no longer visible. This procedure follows the standard practice outlined by Zuur et al. (12). This led to a model allowing for variances to differ between all combinations of levels of species and body segment.

Fewer effects were of interest for comparisons of mean total axial force, as well as mean lateral force ($C_{Fy}$), coefficiencts leading to linear mixed effects models relating each of these to to species, segment, and their interaction. As before, individual was included as a random effect (12, 13), and heterogeneity of variances were handled by introducing weight structures chosen through AIC scoring (12, 14). For total $C_{Fx}$, variance was allowed to differ between combinations of species and segment. For $C_{Fy}$, variance was allowed to differ between body segments.

Efficiency models only had one fixed effect: species. Again, individual was included as a random effect (12, 13), but since variances were equal across species, no weight structure was needed.

Once appropriate models had been fit, ANOVA tests and post-hoc pairwise comparisons could be conducted to determine which effects significantly affected force coefficients. A false discovery rate correction was applied to all post-hoc results to correct for the likelihood that random, false significant differences would be detected (15).

All statistics were performed in R (version 3.5.1, R Foundation for Statistical Computing, Vienna, Austria; https://www.r-project.org/) using the nlme package (version 3.1-137, https://CRAN.R-project.org/package=nlme), and marginal means were estimated for pairwise comparisons using the emmeans package (version 1.2.3, https://CRAN.R-project.org/package=emmeans).

**Data availability**

Raw data, scripts, and extended statistical reports are available to reviewers here https://tufts.box.com/s/sl67axppjkwxb6vcvsjohnmftk668ajp.

Data will be made available to the public upon manuscript acceptance to an accessioned database, e.g., Harvard Dataverse (https://dataverse.harvard.edu/). All scripts used for data analysis will be made available at https://github.com/kelseynlucas.

### SI Figures

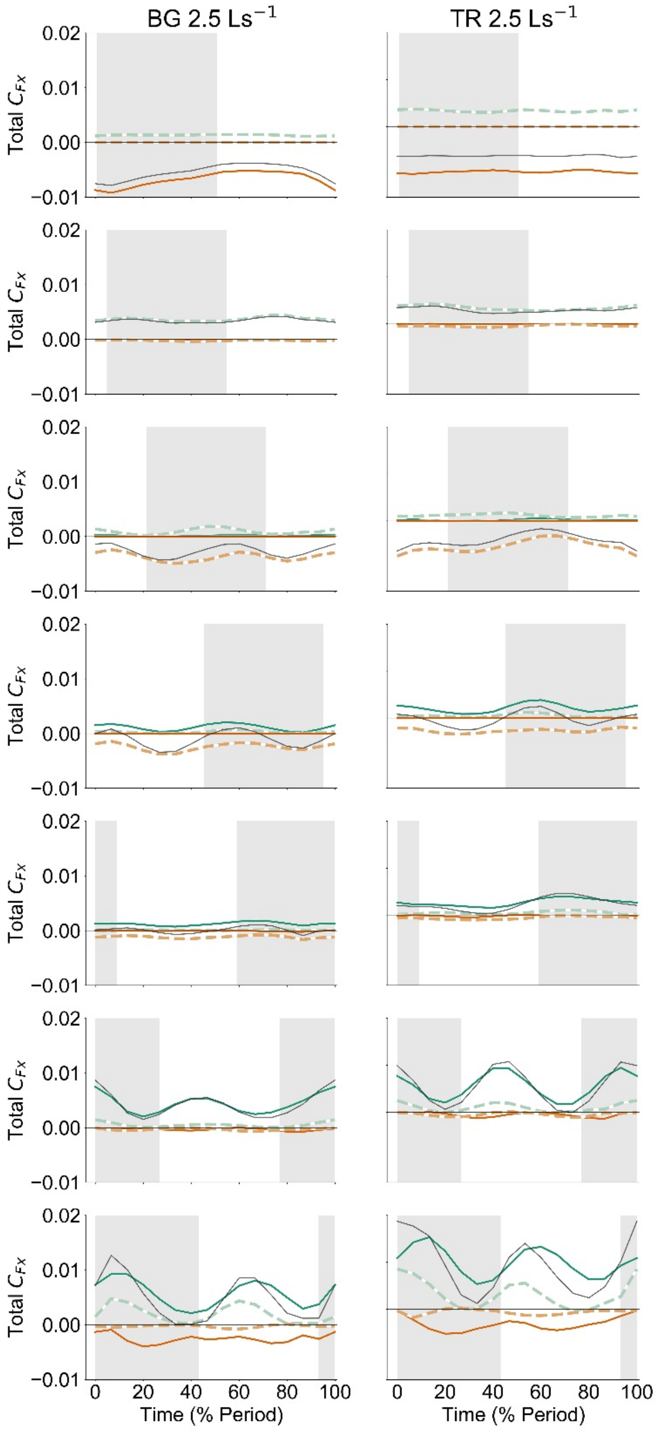


**Fig. S1. Thrust and drag arise from both positive and negative pressure in time- and space-dependent patterns.** Differences in timing of positive and negative pressure thrust forces on the caudal fin (segment 7, bottom row) control the timing of the peak in net thrust. The left column shows the total instantaneous force coefficients acting on the body (contrast with force coefficients from one side of the body in the main text) acting on bluegill (left column) and trout (right column). Rows represent different body segments (see main text). The shaded region in the background indicates the times when the body segment moved from left to right, from peak amplitude to peak amplitude. Forces are colored by force type (green – thrust, orange – drag, grey – lateral component only) and pressure type (dark colors, solid lines – positive pressure, light colors, dashed lines – negative pressure), matching colors in the main text.

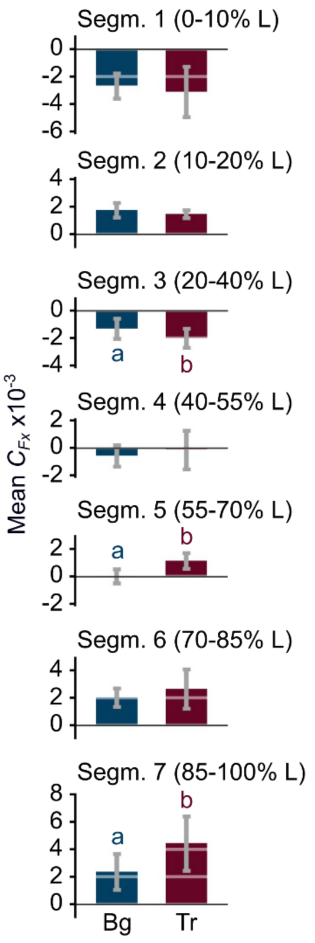


**Fig. S2. Mean streamwise force coefficients in bluegill (Bg) and trout (Tr).** Bars represent time-averaged mean forces on each body segment (Segm.) acting on one side of the body. Forces marked with different letters are significantly different from one another ($p < 0.05$). Transition from net drag to net thrust production occurs on the midbody, but in different segments for each species. The anterior body is net drag-producing, but thrust forces in Segm. 2 greatly reduce the impact of anterior-body drag.

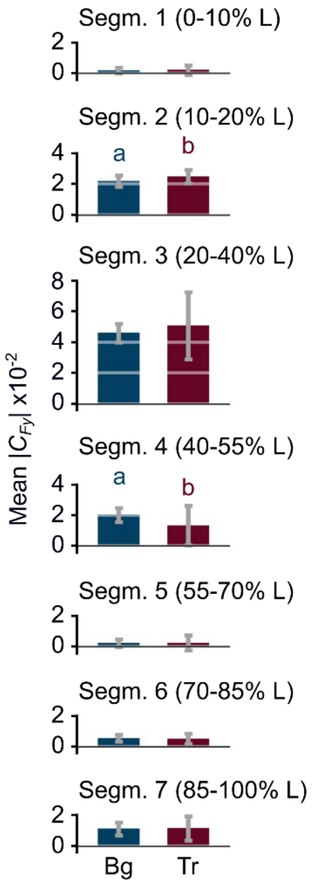


**Fig. S3. Mean lateral force coefficients are similar between bluegill (Bg) and trout (Tr).** The biggest difference occurs in segment 4, where trout have much lower lateral forces than bluegill. Rows represent different body segments (Segm., see main text). Forces marked with different letters are significantly different from one another ($p < 0.05$).

#### SI Movie Captions

**Movie S1. Bluegill velocity field.**

**Movie S2. Trout velocity field.**

**Movie S3. Pressure distribution around bluegill.**

**Movie S4. Pressure distribution around trout.**

**Movie S5. Forces acting on the body of a freely-swimming bluegill.** Forces are colored by force type (green – thrust, orange – drag, grey – lateral component only) and pressure type (dark colors – positive pressure, light colors – negative pressure), matching colors in the main text.

**Movie S6. Forces acting on the body of a freely-swimming bluegill.** Forces are colored by force type (green – thrust, orange – drag, grey – lateral component only) and pressure type (dark colors – positive pressure, light colors – negative pressure), matching colors in the main text.
