## Supplementary figures and images for "The fish body functions as an airfoil: surface pressures generate thrust during carangiform locomotion"

### Supplemental movie 1

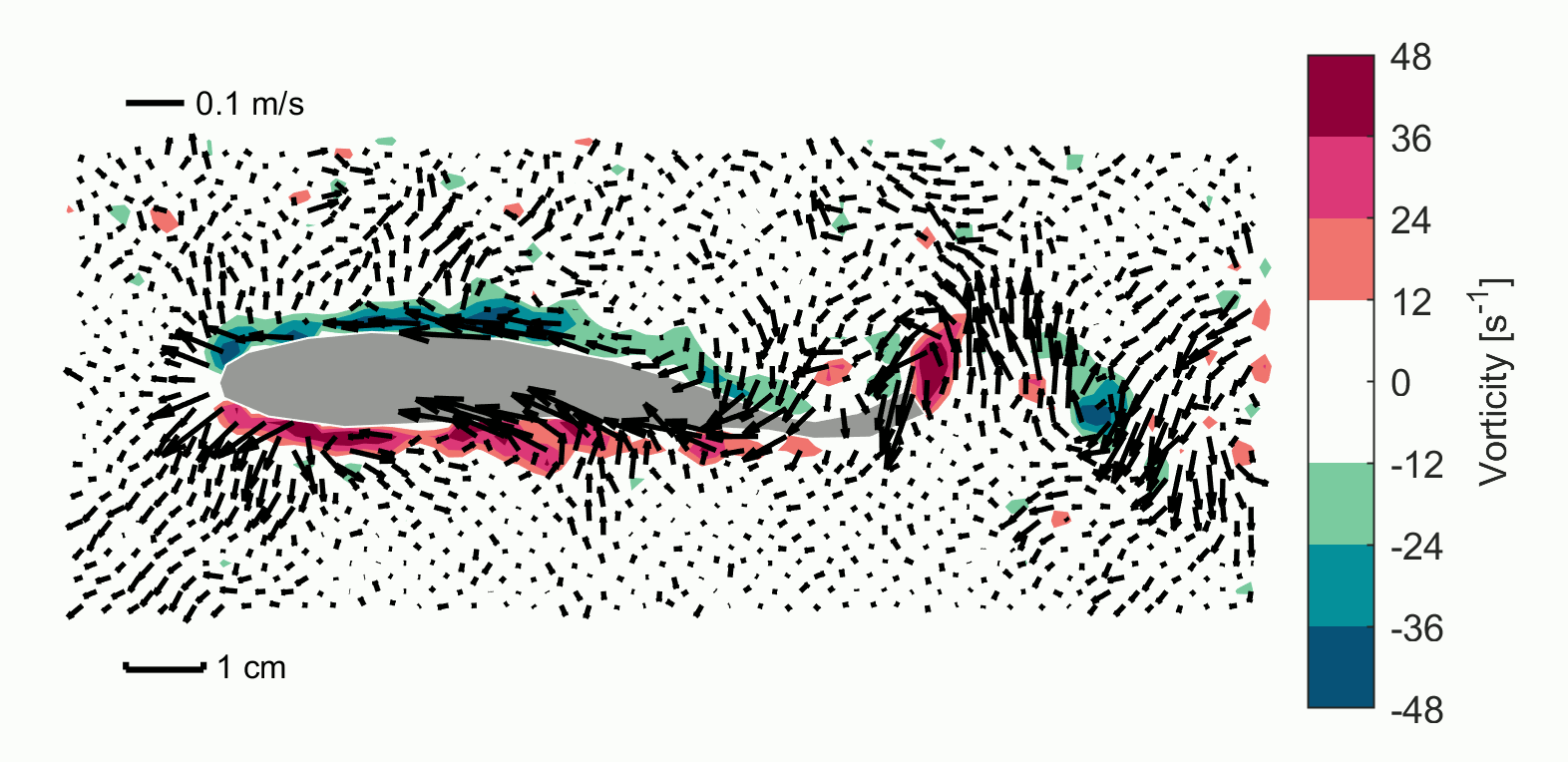

### Supplemental movie 2

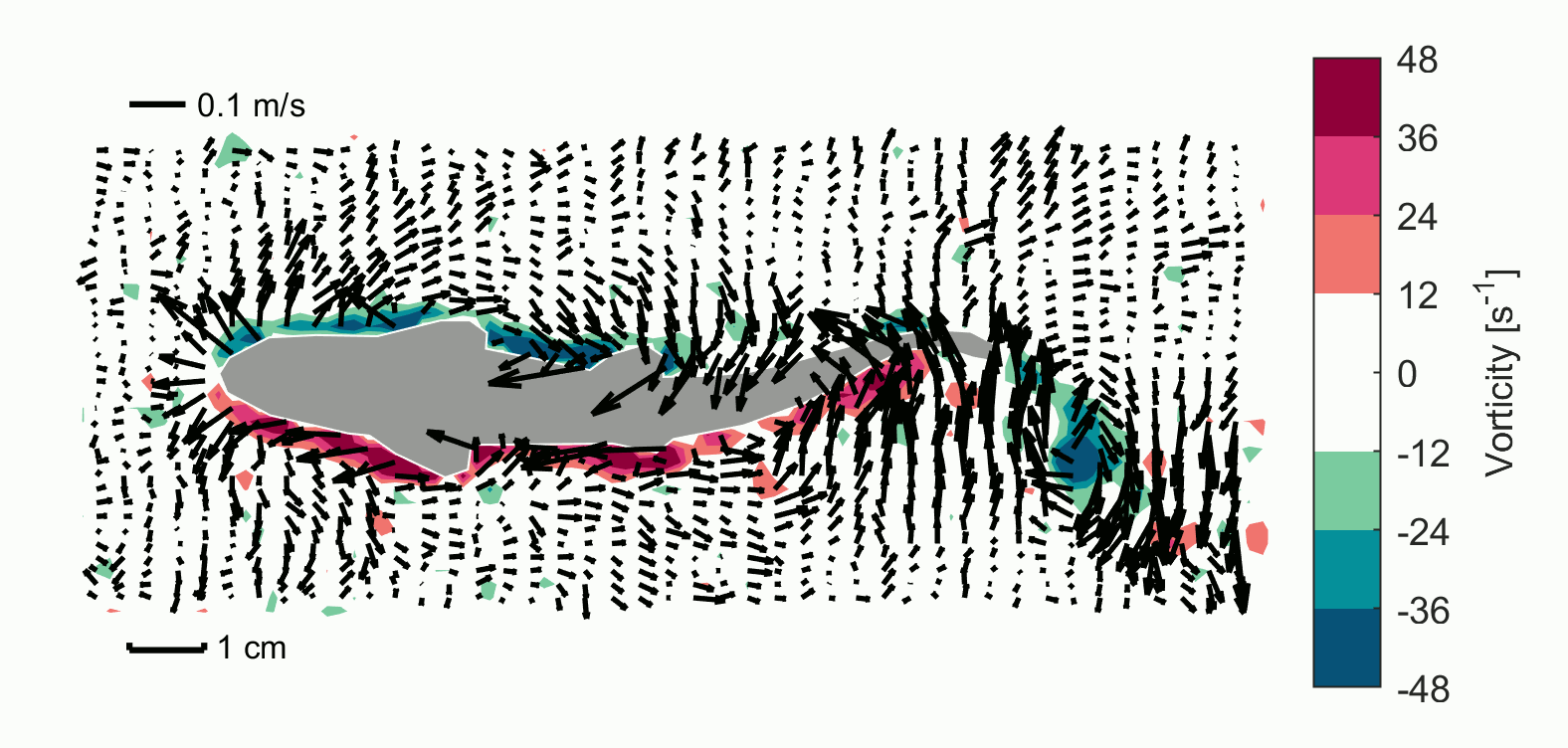

### Supplemental movie 3

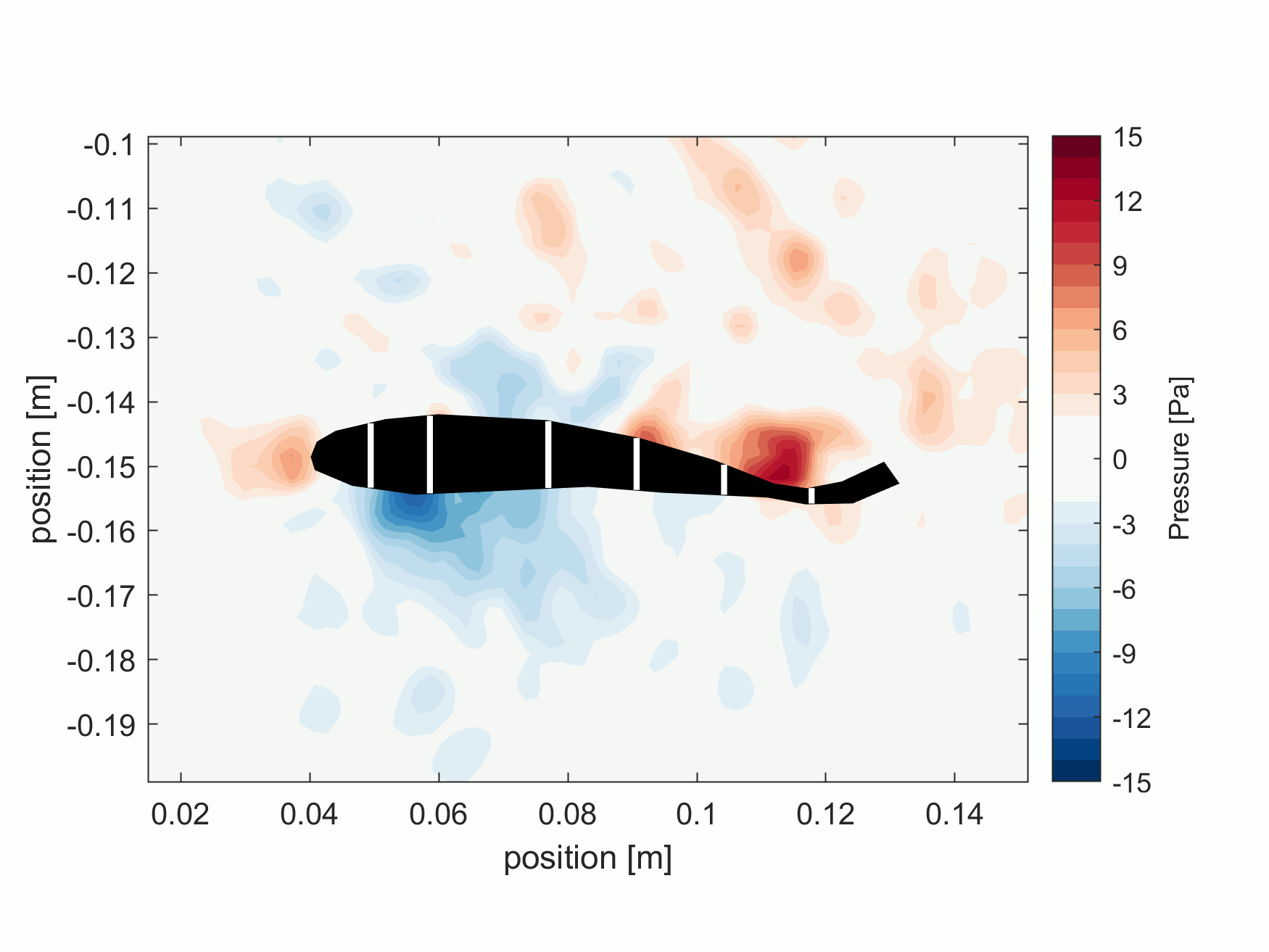

### Supplemental movie 4

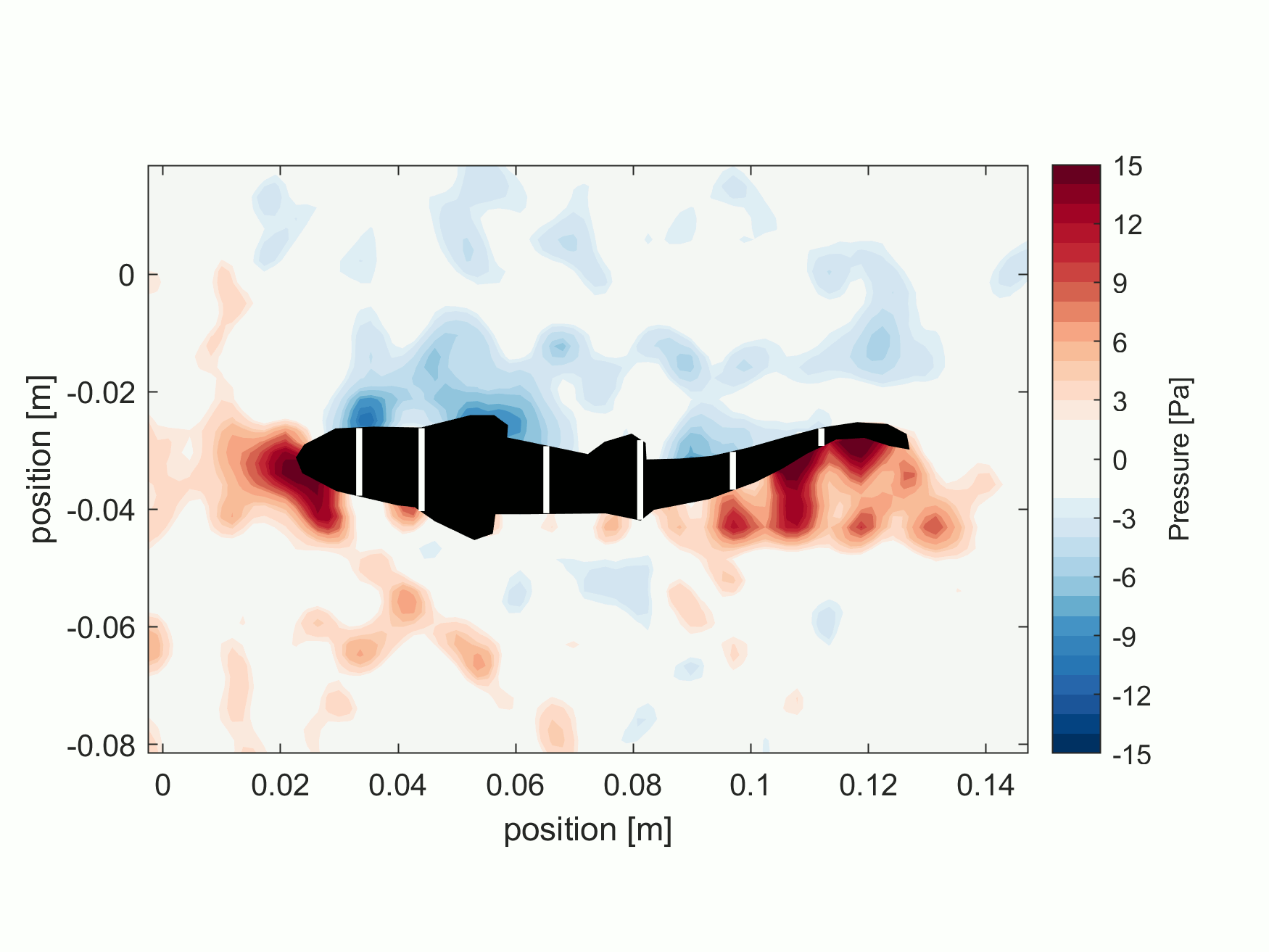

### Supplemental movie 5

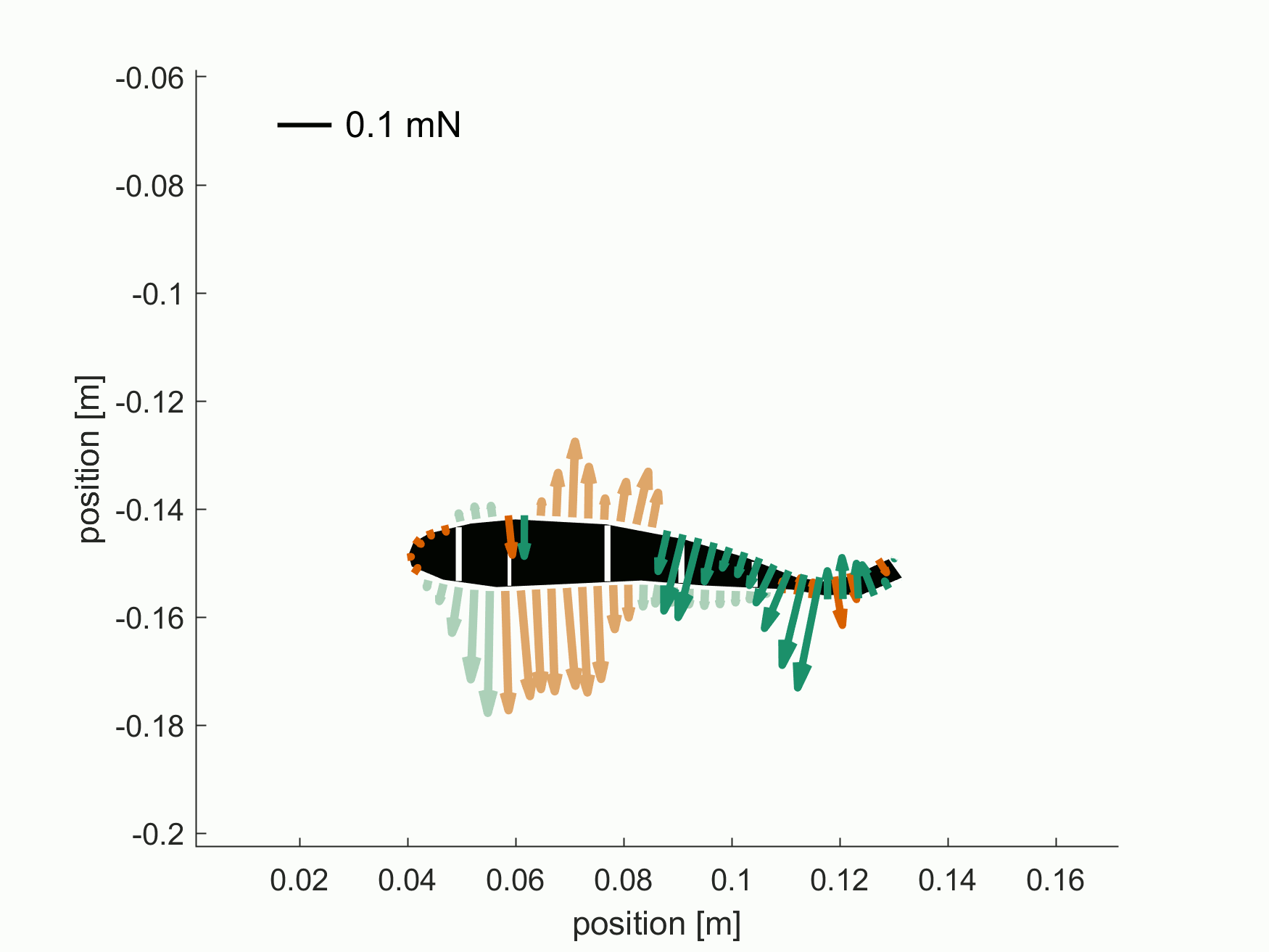

### Supplemental movie 6

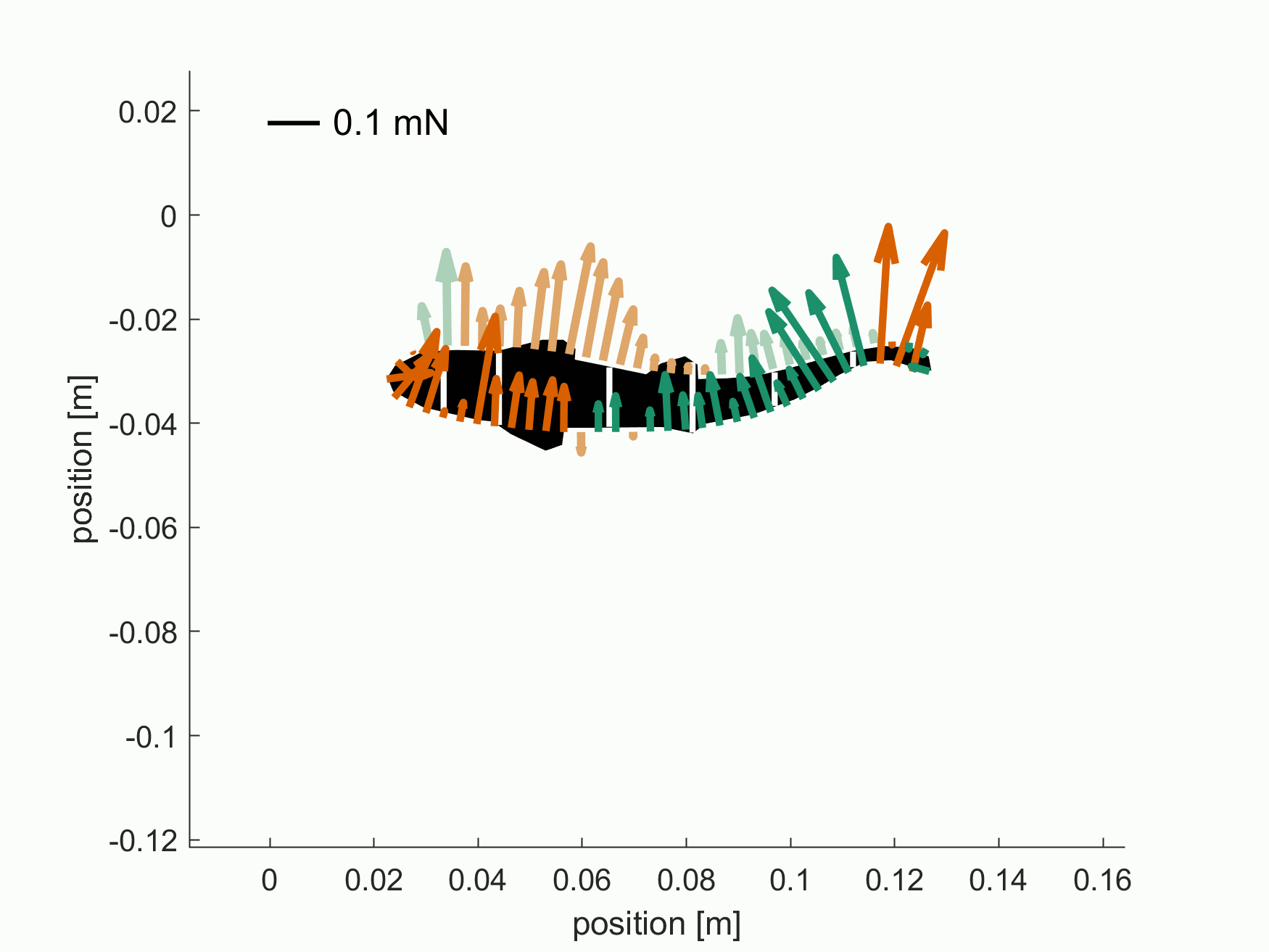
